# Chemical characterization; Antimicrobial and Larvicidal activity of essential oil from *Callistemon citrinus* (Bottle brush) leaves

**DOI:** 10.1101/2021.02.18.431822

**Authors:** D. P. Chachad, A. Dias, K. Uniyal, U. Varma, P. Jadhav, T. Satvekar, M. Ghag – Sawant, M. Mondal, N. Doshi

## Abstract

Mosquito-borne diseases are prevalent in more than 100 countries across the world. They are the major vectors for transmission of Malaria, dengue, yellow fever, filariasis, schistosomiasis, Japanese encephalitis etc. Many of the formerly employed insecticides in mosquito control have harmful effects on human health and other non-target populations, their non-biodegradable nature, the higher rate of bio-magnification in our ecosystem and increasing insecticide resistance on global scale are raising serious concerns. . Therefore, search for natural, eco-friendly alternatives such as bio-insecticides is imperative. In this study, Larvicidal activity of the essential oil obtained from the leaves of *Callistemon citrinus* was tested on Dengue vector mosquito *Aedes aegypti* & Chikungunya vector mosquito *Culex* sp. Also, the chemical composition of the essential oil was recorded using GC-MS analysis. The anti-microbial activity of the essential oil was checked against a few common bacteria and fungi. *Callistemon citrinus* comes out to be one of such bio-insecticides with many therapeutic active constituents, showing appreciable anti-microbial activity and 80-100% larvicidal activity.

## Introduction

Mosquitoes can transmit more diseases than any other group of arthropods and affect millions of people throughout the world. WHO has declared mosquitoes as ‘public enemy no.1’? Mosquito-borne diseases are prevalent in more than 100 countries across the world, infecting over 70, 00, 00, 000 people every year globally and 4, 00, 00, 000 of Indian population. (Ghosh *et. al*, 2012). They are the major vectors for transmission of Malaria, dengue, yellow fever, filariasis, schistosomiasis, Japanese encephalitis etc. Thus, mosquito control has become one of the dire need of the hour which can be achieved only by preventing their proliferation and improving the general cleaning and sanitization protocols for public health. Currently this is being achieved by means of application of synthetic insecticides such as organochloride and organophosphate compounds. (Ghosh *et. al*, 2012) Many of the former synthetic insecticides in mosquito control have been limited due to their harmful effects on human health and other non-target populations, their non-biodegradable nature, the higher rate of bio-magnification in our ecosystem and increasing insecticide resistance on global scale. Therefore, the application of natural, eco-friendly alternatives such as bio-insecticides are imperative for continued effective vector control management.

There is currently a great deal of interest in alternative methods and selective principles for the control of mosquitoes with less environmental damage and which are target specific. In this view, substances from natural origin have immense advantages; they are obtained from renewable resources and the selection of resistant form occurs at slower rate than synthetic insecticides. Another advantage is that they show least or no toxicity to mammals and bees, (deMello, 2017). Shallan *et* al. in 2005, reviewed the current state of knowledge on larvicidal plant species, their extraction processes, growth and reproduction, inhibiting phytochemicals, botanical ovicides, synergistic, additive and antagonistic effects.

*Callistemon citrinus* (Curtis) Skeels is one of such plants, not mentioned in the review, also promises great larvicidal activity. *Callistemon citrinus* (Family: Myrtaceae) commonly known as bottle brush, is frequently cultivated throughout India in gardens as ornamental plant. A handsome shrub or small tree, up to 7.5 m. in height, indigenous to Queensland and New South Wales, is frequently cultivated throughout India in gardens. Leaves are lanceolate, up to 7.5 cm long, with prominent vein, midrib and oil glands; flowers, crimson with dark red anthers, in 10 cm long spikes; capsules depressed-globose. The obvious parts of the flower masses are stamens, mostly red with the pollen at the tip of the filament; the petals are inconspicuous. The essential oils from leaves possess antimicrobial, fungitoxic, antinociceptive and anti-inflammatory activities (Kumar *et. al*., 2011). *Callistemon* species are used for forestry, essential oil production, farm tree/windbreak plantings, degraded-land reclaimation and ornamental horticulture, among other applications (Spencer and Lumley, 1991). In China, *Callistemon* species, especially *C. viminalis*, are used in Traditional Chinese Medicine pills for treating hemorrhoids (Oyedeji *et. al*, 2009). *Callistemon* are also used as weed control (Wheeler, 2005) and as bioindicators for environmental management (Burchett *et. al.*, 2002).

Larviciding is a successful way of reducing mosquito densities in their breeding places before they emerge into adults. In this study, Larvicidal activity of the essential oil was tested on Dengue vector mosquito *Aedes aegypti* & Chikungunya vector mosquito *Culex* sp. Also, the chemical composition of the essential oil was recorded using GC-MS analysis. The anti-microbial activity of the essential oil was checked against a few common bacteria and fungi.

## Materials and Methods

The leaves of *Callistemon citrinus* were collected from Jijamata Udyan (Byculla), Mumbai and were authenticated from R. D. National College Herbarium (RDNCP). They were washed and shade dried for 48 hours and powdered. The essential oil was extracted from 500g of fine powder by hydro distillation using Clevenger’s apparatus for 3 hours. This volatile oil was collected in eppendorf tubes, stored at 10°C under refrigeration until further analysis.

For the study of anti-microbial activity, the extracted oil from *C. citrinus* was checked against 2 bacterial strains viz. *Staphylococcus aureus* and *Eshcherichia coli*, and 2 fungal strains viz. *Candida albicans* and *C. tropicalis* by Vapour diffusion method (Goni P., Lopez P., 2009). *Eucalyptus* oil was used as a positive control.

For the larvicidal activity, 500 ml glass beakers were filled with 250ml of distilled water. About 25 early 3^rd^ and 4^th^ Instar larvae of *Culex sp*. and *Aedes sp*. were transferred from the standard colonies to the beakers containing distilled water. Essential oil from *C. citrinus* was pipetted and spread gently on the surface of the water in the beakers. The percentage mortality for 0.02 ml, 0.04 ml and 0.06 ml was recorded at the end of 24 hours.

Gas chromatography and Mass spectroscopy of the extracted oil was done at S. H. Kelkar Pvt. Ltd. according to the GC-MS protocol for essential oil analysis.

## Results and Discussion

One of the most effective approaches under the biological control program is to explore the floral diversity and enter the field of using safer insecticides of botanical origin as a simple and sustainable method of mosquito control. Further, unlike conventional insecticides which are based on a single active ingredient, plant derived insecticides comprise a blend of phytochemical compounds which act concertedly on both behavioral and physiological processes of the target organisms. (Ghosh, 2012).

The percentage yield of essential oil obtained from the leaves of *C. citrinus* was 1%. Anti-microbial activity of the essential oil showed satisfactory results against test organisms (table no. 1). The larvae were tested with concentration of *C. citrinus* oil of 0.04 ml and 0.06 ml which gave 80-100% mortality rate (table no. 2). GC-MS analysis of the extracted essential oil was carried out and (table no. 3)

**Table no. 1.**

| Essential Oil<br>(0.5 ml) | Antimicrobial Activity on Microbes |  |  |  |
| --- | --- | --- | --- | --- |
|  | *C. albicans* | *C. tropicalis* | *S. aureus* | *E. coli* |
| *Callistemon sp.* | Positive | Positive | Positive | Positive |
| *Eucalyptus sp.* | Negative | Negative | Negative | Negative |

**Table no. 2.**

Table no. 1
| Type of Mosquito Larva | Dose of *Callistemon* oil (ml) | % mortality |
| --- | --- | --- |
| *Aedes* | 0.02 | 20% |
|  | 0.04 | 80% |
|  | 0.06 | 90% |
| *Culex* | 0.02 | 60% |
|  | 0.04 | 100% |
|  | 0.06 | 100% |

**Table no. 3.**
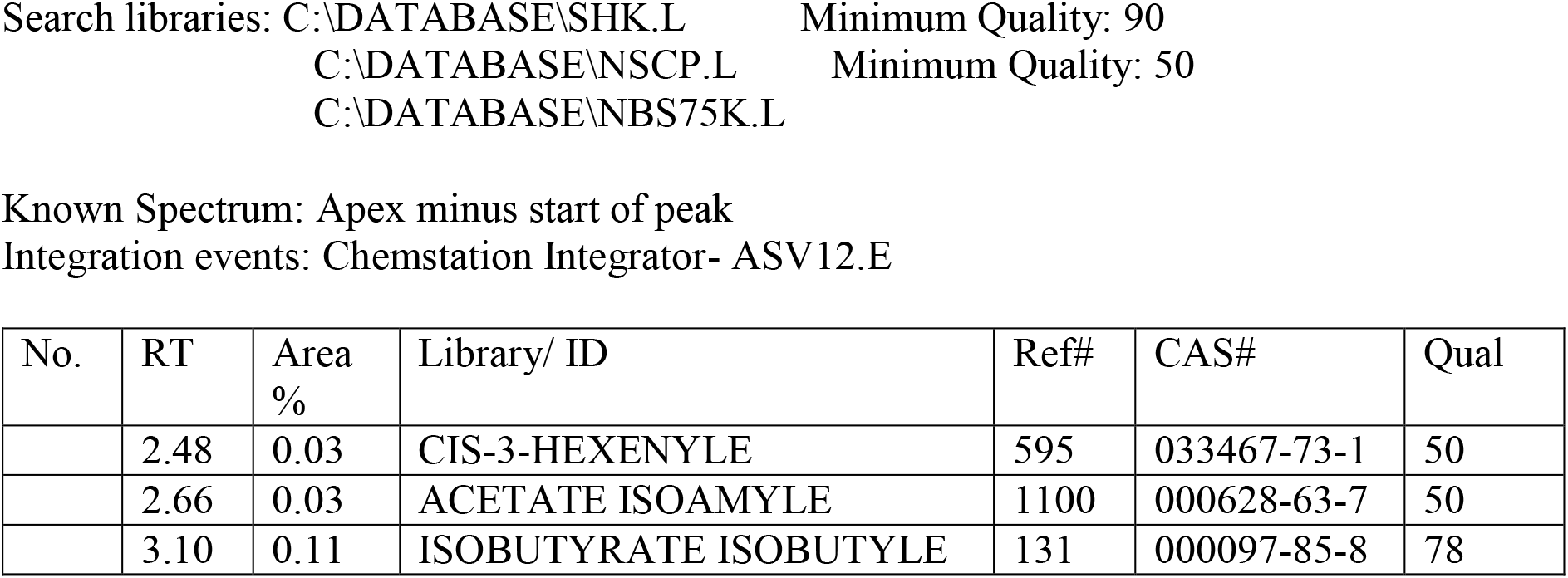
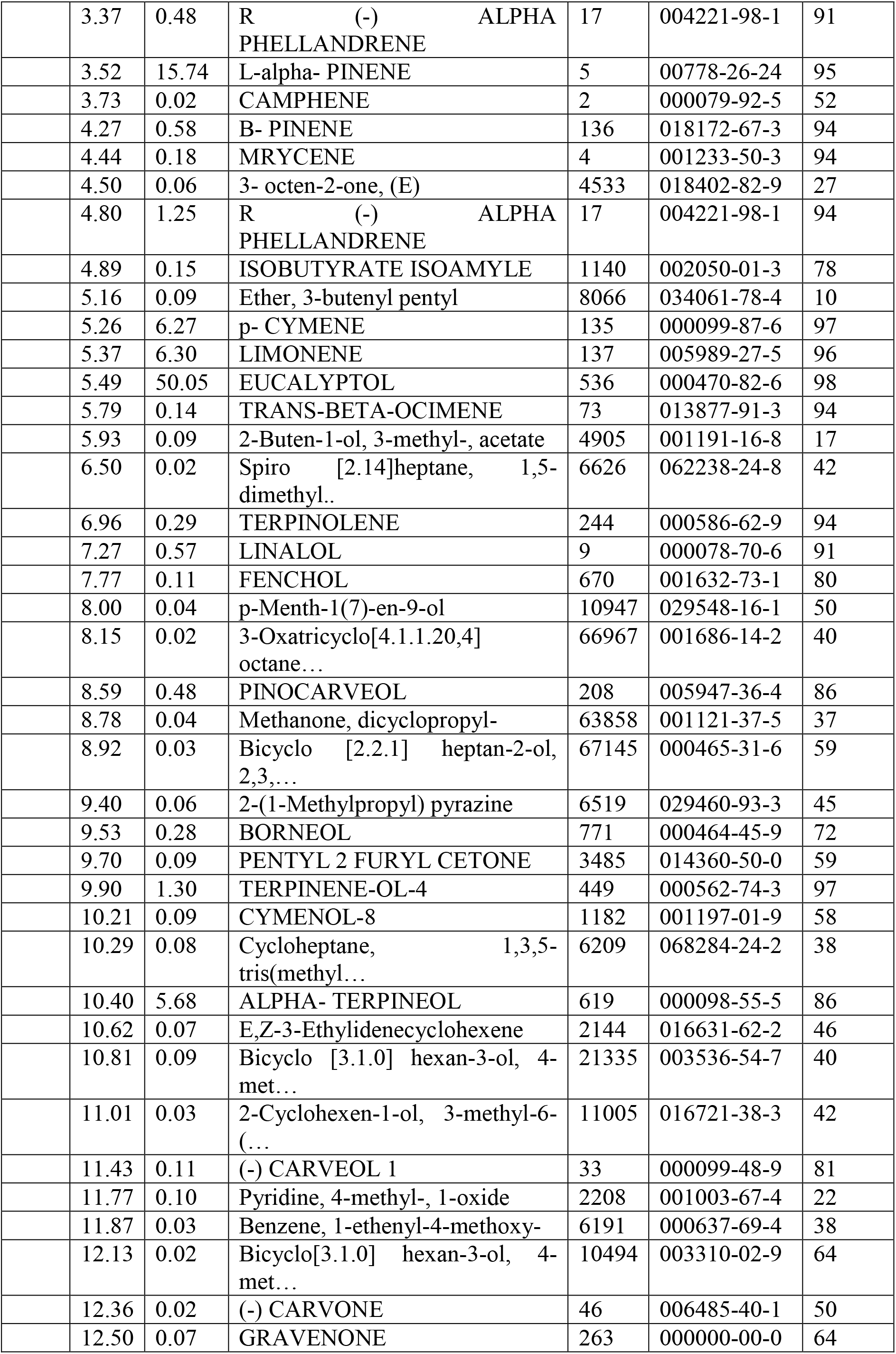
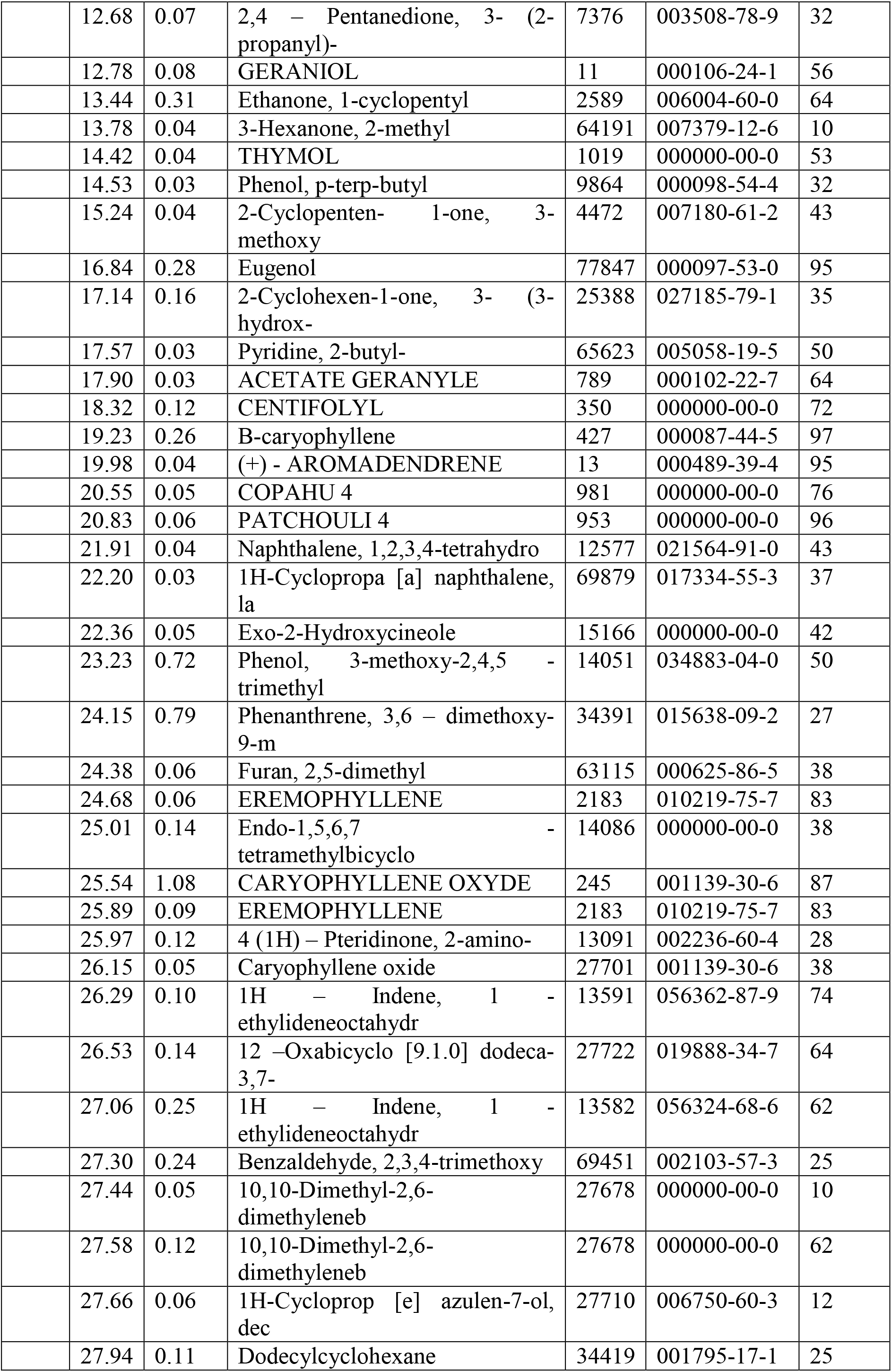
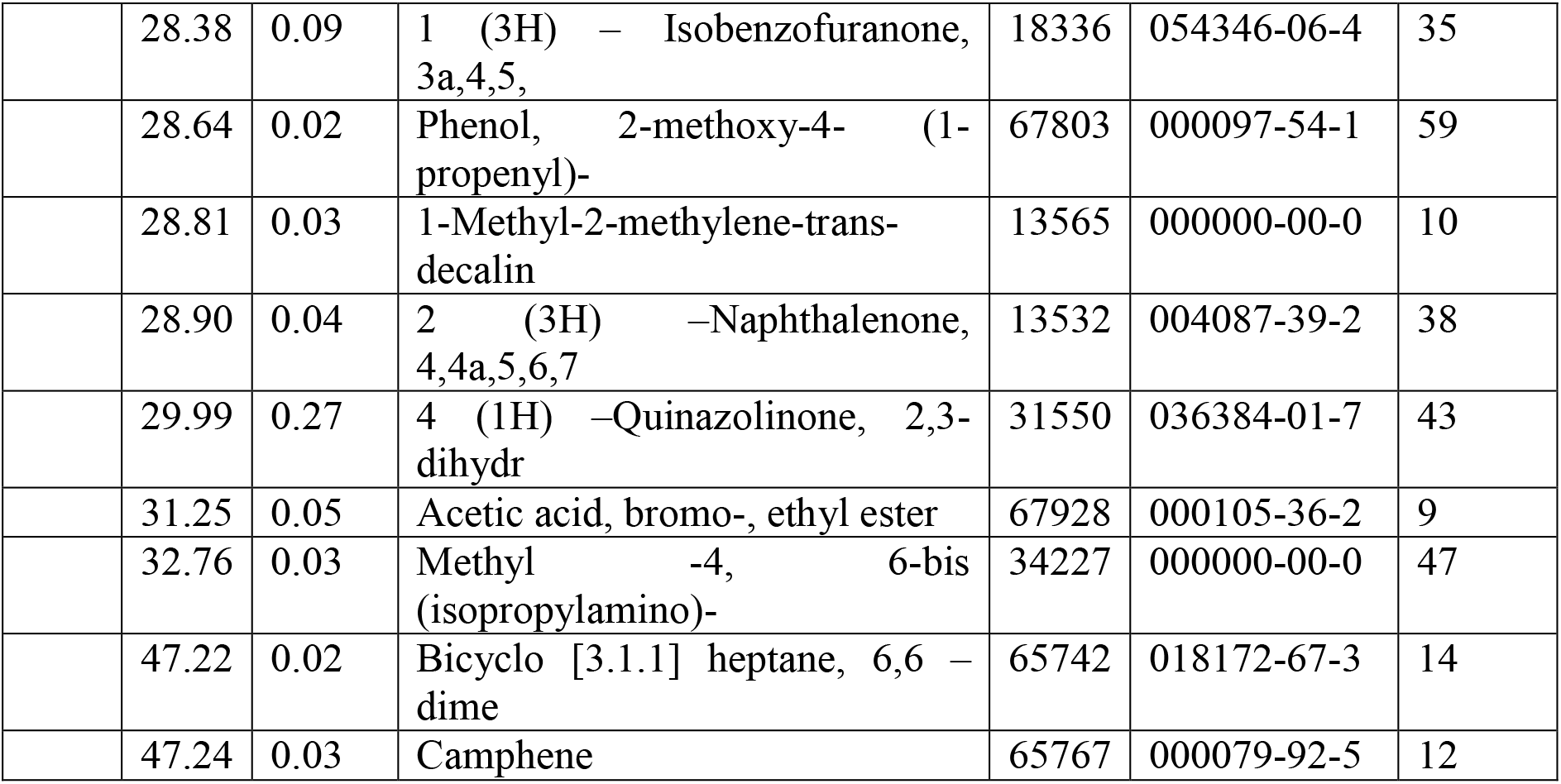

## Conclusion

As the extracted essential oil showed 80-100% mortality against the larvae suggesting that it can be used as a natural larvicidal and can be further tested for insecticidal activity. Positive anti-microbial activity further supports the usage of this essential oil as a surface spray or diluent. Thus, this study takes us one step ahead in the process of bringing natural alternatives for effective mosquito management and larval control for public health.

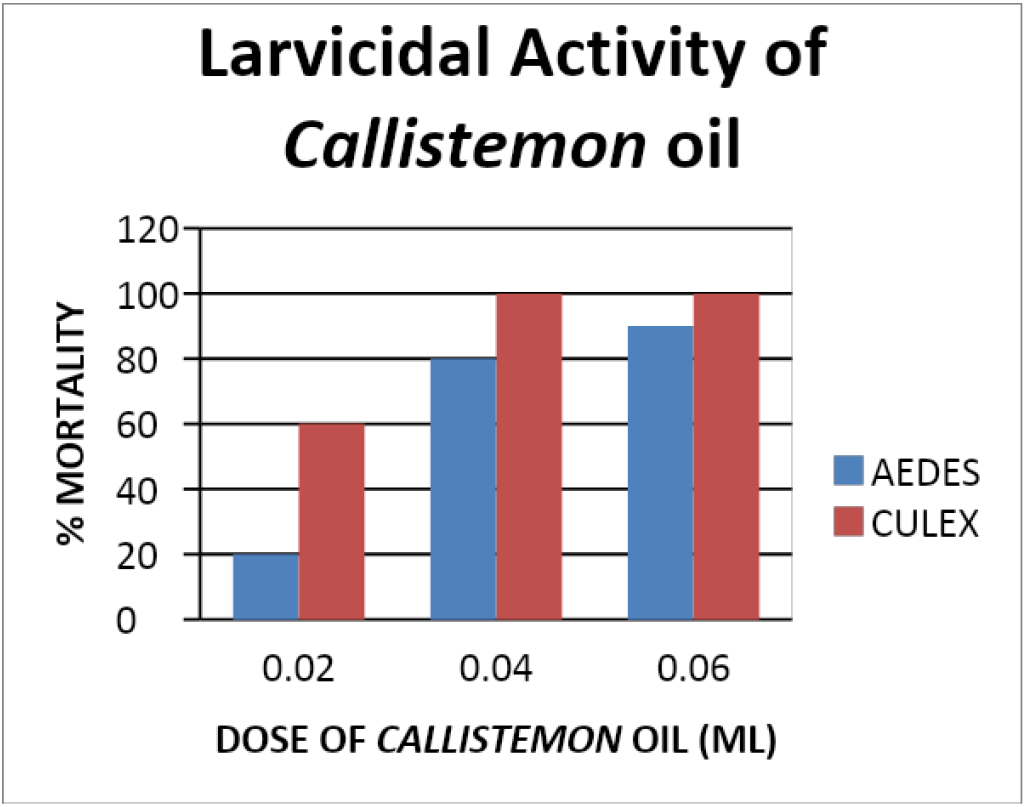

## References

1. Burchett M, Mousine R, Tarran J (2002). Phytomonitoring for urban environmental management. Air Pollut. Plant Biotechnol. 1: 61–91.

2. Das S., Singh S., (2012), Therapeutic potentials of *Callistemon lanceolatus* DC., International journal of advances in Pharmacy, Biology and Chemistry, IJAPBC – Vol. 1(2), Apr-Jun, 2012 ISSN: 2277–4688.

3. de Mello João Carlos Palazzo, Letícia Maria Krzyzaniak, Tânia Mara Antonelli-Ushirobira, Gean Panizzon, Ana Luiza Sereia, José Roberto Pinto de Souza, João Antonio Cyrino Zequi, Cláudio Roberto Novello, Gisely Cristiny Lopes, Daniela Cristina de Medeiros, Denise Brentan Silva, Eneri Vieira de Souza Leite-Mello, 2017, Larvicidal Activity against Aedes aegypti and Chemical Characterization of the Inflorescences of *Tagetes patula*, Evidence-Based Complementary and Alternative Medicine, vol. 2017, Article ID 9602368, 8 pages.

4. Ghosh Anupam, Chowdhury Nandita, Chandra Goutam (2012), Plant extracts as potential mosquito larvicides, Indian Journal of Medical Research, Year 2012, volume 135, issue 5, page 581–589.

5. Kumar, Sunil & Kumar, Virat & Prakash, om. (2011). Pharmacognostic study and anti– inflammatory activity of *Callistemon lanceolatus* leaf. Asian Pacific journal of tropical biomedicine. 1. 177–81. 10.1016/S2221-1691(11)60022-1.

6. Oyedeji, O. O., Lawal, O. A., Shode, F. O., & Oyedeji, A. O. (2009). Chemical composition and antibacterial activity of the essential oils of *Callistemon citrinus* and *Callistemon viminalis* from South Africa. Molecules (Basel, Switzerland), 14(6), 1990–1998.

7. Shaalan, Essam & Canyon, Deon & Younes, Mohamed & Abdel-Wahab, Hoda & Mansour, Abdel-Hamid. (2005). A review of botanical phytochemicals with mosquitocidal potential. Environment international. 31. 1149–66. 10.1016/j.envint.2005.03.003.

8. Spencer RD, Lumley PF (1991). *Callistemon*: In Flora of New South Wales; Harden GJ, Ed.; New South Wales, University Press: Sydney, Australia., 2: pp. 168–173.

9. The Wealth of India (1992). A Dictionary of Indian Raw Materials and Industrial Products, Vol-3: Ca-Ci, Publication and information Directorate, Council of Scientific and Industrial Research, New Delhi.

10. Wheeler, Greg. (2005). Maintenance of a narrow host range by *Oxyops vitiosa*; a biological control agent of *Melaleuca quinquenervia*. Biochemical Systematics and Ecology. 33. 365–383. 10.1016/j.bse.2004.10.010.

